# Magnetic entropy as a gating mechanism for magnetogenetic ion channels

**DOI:** 10.1101/148379

**Authors:** Guillaume Duret, Sruthi Polali, Erin D. Anderson, A. Martin Bell, Constantine N. Tzouanas, Benjamin W. Avants, Jacob T. Robinson

## Abstract

Magnetically sensitive ion channels would allow researchers to better study how specific brain cells affect behavior in freely moving animals; however, recent reports of “magnetogenetic” ion channels have been questioned because known biophysical mechanisms cannot explain experimental observations. Here we show that magnetic fields can produce a change in the magnetic entropy of biogenic nanoparticles, which in turn may generate sufficient heat to gate temperature-sensitive ion channels. This magnetocaloric effect provides a rational approach for developing future magnetogenetic channels.

---

Genetically encoded ion channels that open in response to magnetic fields - “magnetogenetics” – is a potentially powerful technique for neuroscientists. Because magnetic fields can freely penetrate bone and tissue, cells that express these magnetogenetic proteins could be activated or inactivated throughout the brain of freely moving animals without the need for any implanted probes. Compared to other noninvasive brain stimulation techniques like ultrasound ^1^ and transcranial magnetic ^2^ or electric fields ^3,4^, magnetogenetics could target cells of a specific genetic identity. This genetic specificity is one of the main advantages of techniques like opto-^5^, thermo-^6^, and chemo-genetics ^7^. Compared to these existing technologies, magnetogenetics would provide an important complement. Because magnetic fields could be applied independent of optical, and electrical methods to stimulate or record brain activity, scientists could simultaneously probe multiple distinct cell populations in the same organism. Magnetic fields also have distinct advantages as a stimulus. Compared to light, magnetic fields provide superior penetration in biological tissue. Compared to thermogenetics and chemogenetics, magnetogenetics provides superior temporal resolution because it does not rely on heat or chemicals to diffuse through the target tissue.

Typically, genetically targeted brain stimulation techniques are developed by selecting specific genes from organisms that naturally respond to a particular stimulus; however, there are no naturally occurring genes that are known to produce magnetic sensitivity when expressed in a host cell or organism. A major challenge in identifying the genetic basis of natural magnetoreception is the fact that there is no scientific consensus regarding the biophysical mechanism except in the case of magnetotactic bacteria ^8^. Without a natural magnetic field receptor, scientists are forced to engineer synthetic proteins that respond to magnetic fields. One approach is to combine chemically synthesized magnetic nanoparticles that, when injected into an organism, associate with temperature-sensitive ion channels to create magnetically sensitive cells^9,10^. High frequency alternating magnetic fields (typically 100- 500 KHz) can heat these nanoparticles through relaxation losses^11^ and activate the associated thermoreceptor. While these injected nanoparticles effectively create magnetically sensitive cells, a completely genetically encoded strategy would greatly simplify the implementation of magnetogenetics.

Recently, two independent labs have reported synthetic magnetogenetic protein assemblies based on the iron binding protein ferritin ^12,13;^ however, no established biophysical mechanisms can explain the observed magnetic responses ^14,15^. In each case, the reported magnetogenetic constructs were based on tethering genetically encoded iron nanoparticles assembled within a 24-mer ferritin cage to ion channels of the TRP-family ^16^. It has been argued that the heat produced by ferritin in an alternating magnetic field is too weak to activate the nearby thermoreceptors ^14,15^. Surprisingly, several papers also report channel gating by steady or low frequency magnetic field that is not expected to produce heat ^12,13^. This channel gating has been proposed to be mediated by the mechanical force between adjacent nanoparticles, but these forces are at least eight orders of magnitude weaker than the pN-scale forces required to activate mechanoreceptors ^14^. Magnetically induced eddy currents responsible for transcranial magnetic stimulation (TMS) also fail to account for the observed magnetic response because TMS requires voltage gated ion channels ^17^. The TRPV4 channel, which was used in the “*Magneto2.0*” construct ^12^, has negligible voltage sensitivity (Fig. S3).

## Results and discussion

Here, we focus on channel gating by slowly varying magnetic fields and propose that the magnetocaloric effect - a phenomenon that, to our knowledge, has not been reported in biological systems - explains the previously reported magnetogenetic channel activation.

The magnetocaloric effect is based on a conversion between magnetic entropy and heat. In the absence of a magnetic field, the ensemble of magnetic moments within a paramagnetic material are randomly oriented, yielding no net magnetic moment (Fig. 1a). Similarly, in superparamagnetic materials a single magnetic domain rapidly reorients such that the particle displays no net magnetic moment when measured over periods of time longer than the relaxation time^18^. For small particles like ferritin this relaxation time is expected to be on the order of nanoseconds^19^. In either case of paramagnetic or superparamagnetic materials, when a magnetic field is applied, the moments align while the field is present, thereby reducing the magnetic entropy (Fig. 1a).

**Fig. 1.**
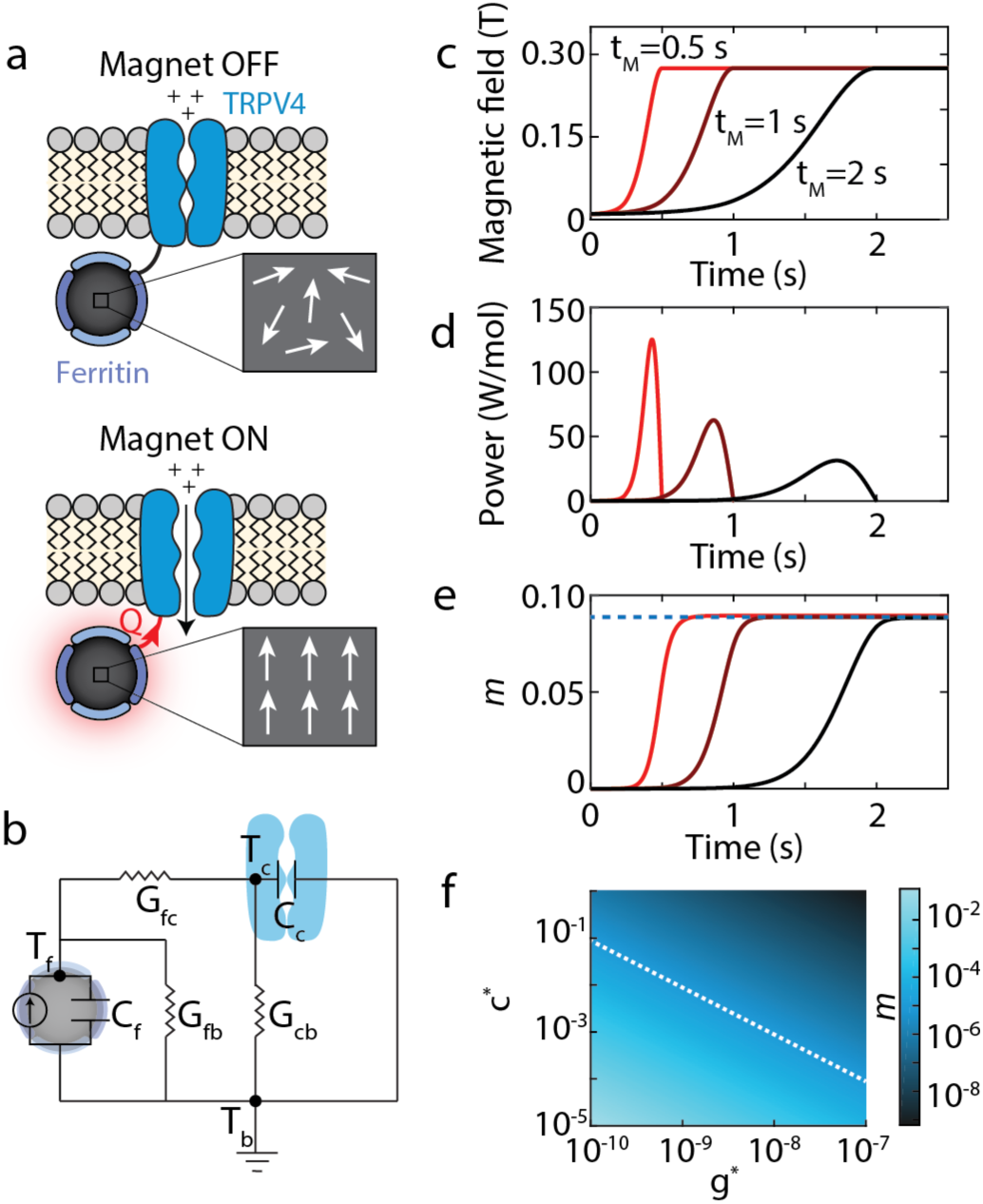
The magnetocaloric gating mechanism: (**a**) Schematic shows how the magnetocaloric effect in ferritin can activate nearby temperature-sensitive ion channels (e.g. TRPV4). An applied magnetic field will align the magnetic moments within paramagnetic ferritin nanoparticles, which will reduce the magnetic entropy. The reduced magnetic entropy generates heat (*Q*) via the magnetocaloric effect that can activate a nearby temperature-sensitive ion channel. Here we have depicted ferritin as a paramagnet, but the calculations are equivalent for superparamagnetic particles. (**b**) Equivalent circuit model used to estimate heat transfer between the ferritin particle and ion channel. *T_f_*, *T_c_*, *T_b_* represent the temperature of the ferritin, channel and bath, respectively. *C_f_* and *C_c_* represent the heat capacities of the ferritin and channel, respectively.G_fc,_ G_fb_, G_cb_ represents the thermal conductances between the ferritin and channel, ferritin and bath, and channel and bath respectively. (**c**) Applied magnetic field as a function of time for three different magnetization times (*t_m_*= 0.5 s, 1 s, and 1.5 s). (**d**) The power generated in a mole of ferritin particles due to magnetocaloric effect for the magnetic field profiles in c. (**e**) Fraction of channels (m) gated by the magnetocaloric effect based on (d) and Eq. 2.The dashed blue line indicates the maximum percentage of channels that open as derived by the analytical expression for *m* in Eq. 3. Note that the total number of channels that open depends on the maximum value of the magnetic field and not the rate of magnetization. Calculations assume *T_b_* = 25°C, *c*\* = 10^−5^ and *g*\* = 10^−10^ (**f**) The fraction of channels that respond depends on the value of *c*\* (heat capacity scaling factor) and *g*\* (thermal conductance scaling factor), which can vary by orders of magnitude depending on the biophysical mechanism that triggers temperature-dependent channel gating. Dashed white line corresponds to the range of values for c* and g* that give m = 10^−5^. We expect that the m values near this limit and greater would yield a physiological response.

This decrease in magnetic entropy is compensated by an increase in molecular vibrations which produces heat (*Q*) ^20^.

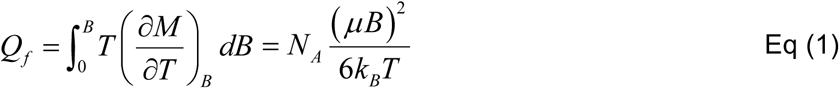

Here *M* is the magnetization of ferritin, which we approximate using the Langevin function, *B* is the applied field and *T* is the temperature of the bath. Evaluating this integral gives us an expression in terms of the magnetic moment of ferritin nanoparticles, which is known to depend on the number of iron atoms in the nanoparticle as well as the mineralization/oxidation of the iron core ^21^. For our calculations, we assume a magnetic moment of μ =316μ_B_, where μ_B_ is the Bohr Magneton. This value of magnetic moment corresponds to approximately 4000 iron atoms ^19^ (less than the 4500 iron atoms reported for fully loaded ferritin^22^). Based on Eq. 1 we calculate that a 275 mT magnetic field will generate 15.8 J/mol of ferritin. (Supplementary Information (SI) Section 1.1). While most literature suggests that ferritin is superparamagnetic ^19,23^, this calculation is identical for superparamagnetic or paramagnetic materials. Using an equivalent circuit model (Fig. 1b) and estimates of the interfacial area between the ferritin and the channel, we expect approximately η=16% of this heat will reach the channel (see SI Section 1.2).

To determine if the roughly 2.5 J/mol that reaches the channel due to the magnetocaloric effect is sufficient to gate TRP thermoreceptors, we can estimate the number of additional channel openings (*m*) based on a temperature-dependent increase in the channel open rate (*a*) and the probability that a channel is in a closed activatable state (*P_c_*):

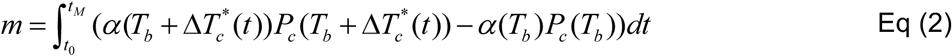

Here *P_c_* is the probability of channel being closed, *α* is the channel opening rate, T_b_ is the bath temperature and *ΔT*\**_c_* is the change in effective temperature of the ion channel (see SI Section 1.3). Because heat is applied only during magnetization, this integral is evaluated only during the time that the magnetic field is changing (t_0_ to t_M_). Note that we are assuming that the magnetization is fast enough to neglect any adaptation by the cell to a change in temperature. This adaptation typically involves transcriptional regulation of calcium pumps (PMCA) and ion exchangers (NCX) time scale of minutes ^24,25^ on the tens of. In our experiments, the magnetization time is less than 1 second.

In principle, it is possible to estimate the local temperature change (*ΔT*) due to the magnetocaloric effect and to evaluate the integral in Eq. 2 to determine the additional channel openings; however, nanoscale temperature changes are not well understood for nanoparticles in solution^26^. Two major factors are believed to make it difficult to estimate the temperature near the surface of magnetic nanoparticles: 1) nanoscale interfacial thermal conductance can be orders of magnitude lower than in macroscopic systems ^27,9,26,28,29^ and 2) steep temperature gradients can cause some regions of the protein to have a higher temperature than would be predicted if the channel was heated uniformly (see SI Section 1.2). Although the exact values of the thermal conductance and effective local temperature is unknown for this protein assembly, we can introduce two factors: *g*\* and *c*\*, where the true thermal conductivity can be written as *g*\**G* (where *G* is the macroscopic thermal conductance), and the effective change in temperature at the critical protein domain can be written as *ΔT/c*\* (where *c*\* is a heat capacity scaling factor) (see SI Section 1.2). Evaluating Eq. 2 then gives us an estimate for *m* given these two parameters (see SI Section 1.3):

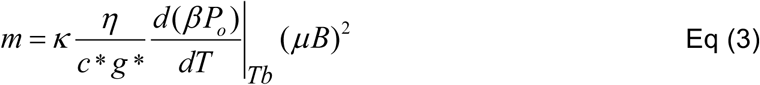

where β is the closing rate of the channel, P_o_ is its open probability and κ is defined as:

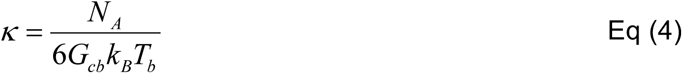

Based on theory and experiments we can then set bounds on the values for *c*\* and *g*\* and find the range of channel openings (*m*) expected from the magnetocaloric effect. For example, multiple experiments suggest that at the nanoscale, heat dissipation rates can be up to 10 orders of magnitude smaller compared to macroscopic systems ^27,9,26,28,29^. Thus, we expect *g*\* to range between 10^−10^ and 1. Similarly, we estimate the scaling factor for heat capacity of TRP channels, *c*\* to be between 10^−5^ and 1 (see SI Section 1.2). The upper bound of the effective heat capacity represents uniform heating of the channel with heat distributed evenly to all degrees of freedom. On the other hand, the lower bound corresponds to all heat being absorbed locally by a single degree of freedom, such as the breaking of a hydrogen bond or the rotation of a protein side chain before the protein reaches thermal equilibrium (see SI Section 1.2). In the case of TRPV4 a single hydrogen bond between residue L596 in the S4-S5 linker and residue W733 in the TRP domain has been proposed as a “latch” that stabilizes the protein in the closed and inactivated state ^30^. The breaking of this hydrogen bond due to increased temperature, which can occur in tens of picoseconds ^31^, is believed to destabilize the protein leading to channel gating. Interestingly this hydrogen bond is estimated to be only about ~3.5nm from the ferritin binding site based on a homology model for *Magneto2.0* ^30^.

Plotting *m* over the range of expected thermal conductivities and effective heat capacities we find that at the extreme, approximately 1 in 10 channels would be activated by applying a 275 mT magnetic field (Fig. 1f). Transfected hippocampal neurons can express between 160,000 and 1,000,000 heterologous functional TRPV1 channels ^32^, thus we would expect magnetic responses that could be as large as approximately 10,000 to 100,000 additional channel openings per cell. TRPV4 channels have a conductance of 60 pS ^33^, and the activation of a single ion channel with conductances of 60-70 pS can trigger action potentials in neocortical and hippocampal neurons ^34^ (see SI Section 1.4). Transfected HEK cells, on the other hand, are expected to express approximately 1000 exogenous ion channels ^35^. Near the maximum value of *m*, we anticipate about 100 magnetically activated channel openings per HEK cell. Given the high conductance of these channels we expect approximately 10^5^ Ca^2+^ ions to enter the cell per channel opening, which is near the minimum detection limit for Fluo-4 (approximately 10^5^ Ca^2+^ ions; see SI Section 1.5). Overall, our model predicts that a combination of low thermal conductance at the nanoscale combined with local heat absorption by the channel protein could produce responses similar to what was reported in Wheeler et al. ^12^ and Stanley et al. ^13^.

To test if the magnetic sensitivity reported for TRP-ferritin fusion proteins is indeed thermally mediated, we designed experiments to selectively inhibit either the thermal or mechanical sensitivity of *Magneto2.0* ^12^. The TRPV4 channel is known to have separate activation pathways for temperature and mechanical stress in HEKs ^36^, allowing us to independently attenuate the mechanical and thermal sensitivity of the channel (Fig. 2a). To reduce the mechanical sensitivity of *Magneto2.0* we used a PLA2 inhibitor, 4-bromophenacyl bromide (pBPB)^36,37^. This condition, referred to as *(-)Mech,* showed reduced sensitivity to hypo-osmotic shock but normal temperature sensitivity as measured by calcium sensitive fluorescence imaging in transfected HEK cells (Fig. 2b, c, *(-)Mech*). Similarly, we created a version of *Magneto2.0* with reduced thermal sensitivity by mutating the YS domain in the third transmembrane domain (Y555A/S556A) ^36^. This variant, referred to as *(-)Therm,* showed normal response to hypo-osmotic shock, but reduced temperature sensitivity (Fig. 2b,c, *(-)Therm*).

**Fig. 2.**
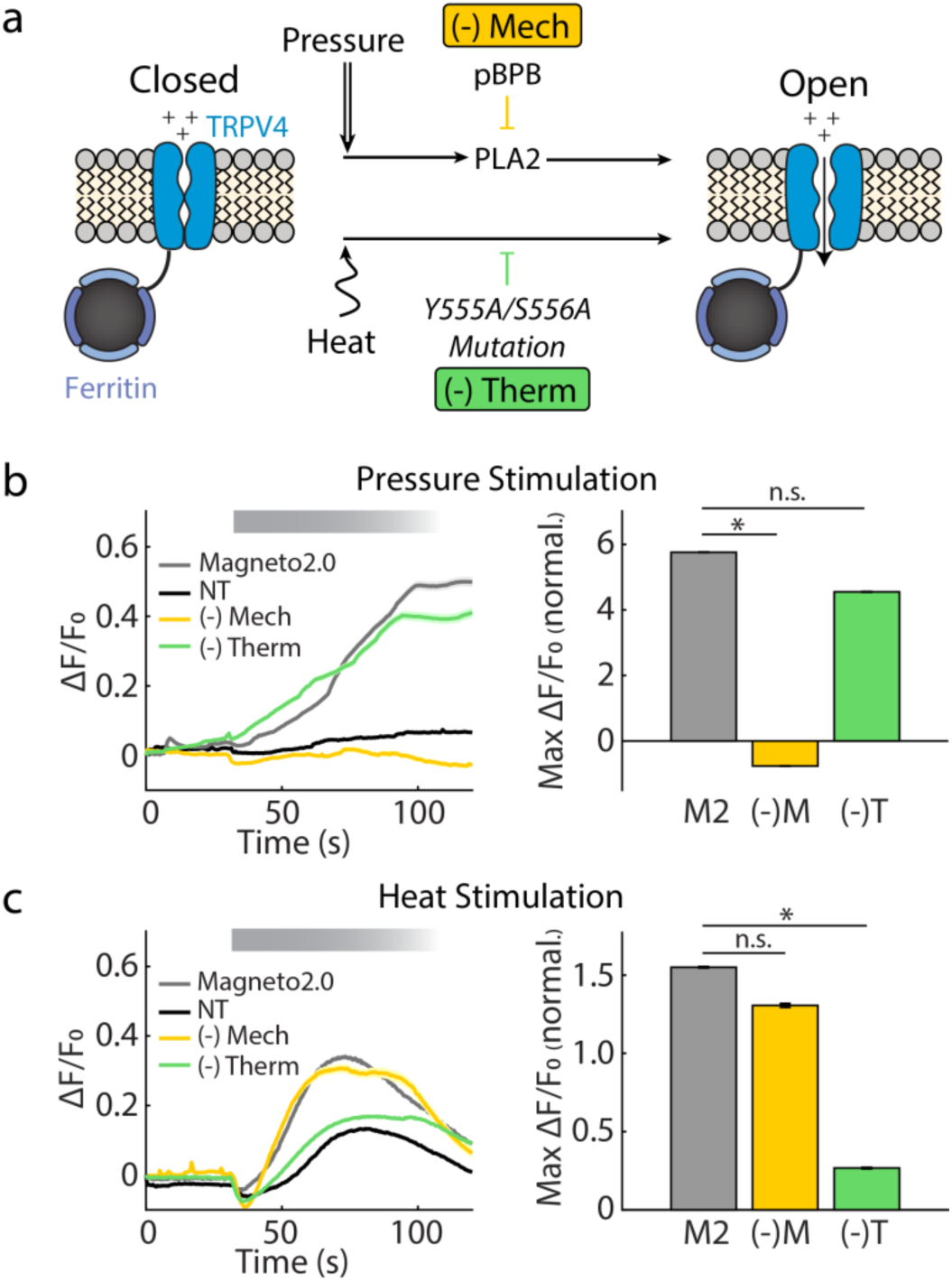
Inhibition of distinct activation pathways in *Magneto2.0*: (**a**) Schematic depicting independent pathways by which TRPV4 responds to stimuli. pBPB inhibits the PLA2-dependent mechanical response of TRPV4 ^36^. We refer to this condition as *(-)Mech*. The mutation Y555A/S556A inhibits the thermal response of TRPV4 ^36^. We refer to this condition as *(-)Therm*. (**b**) Calcium-sensitive fluorescence imaging shows that *(-)Mech* (and not *(-)Therm*) has reduced sensitivity to hyposmotic stimulation compared to WT *Magneto2.0*. (NT, n = 3473 cells from 6 separate cell cultures; *Magneto2.0*, n = 660 cells from 6 separate cell cultures; *(-)Mech*, n = 711 cells from 7 separate cell cultures; *(-)Therm*, n= 338 cells from 5 separate cell cultures). (**c**) Calcium-fluorescence imaging shows that *(-)Therm* has a significantly reduced response to thermal stimulation (40°C perfusion) compared to *(-)Mech* and *Magneto2.0*. (NT, n=1482 cells from 3 separate cell cultures; *Magneto2.0*, n= 395 cells from 5 separate cell cultures; *(-)Mech*, n = 578 cells from 4 separate cell cultures; *(-)Therm*, n = 343 cells from 6 separate cell cultures). Bold lines in ΔF/F_0_ vs. time represent mean values and shaded regions represent ± s.e.m. based on the number of cells recorded. Bar plots represent the maximum ΔF/F_0_ normalized to the maximum ΔF/F_0_ for non-transfected cells (NT). Except for NT, data are obtained only from mCherry^+^ (transfected) cells. The significances are assessed with a two-tailed unpaired Student’s t-test, (*: P < 0.001).

We found that the *(-)Mech* variant of *Magneto2.0* responded to magnetic stimulation while the *(-)Therm* variant did not, suggesting that magnetic sensitivity in slowly varying fields is indeed a thermal response as predicted by our magnetocaloric hypothesis. In WT *Magneto2.0* and *(-)Mech,* we observed a slow increase in intracellular calcium when we applied a magnetic field of approximately 275 mT at a frequency of 0.08 Hz (Fig. 3). We note that this slow increase in calcium is similar to the data reported by Wheeler et al. in transfected HEK cells ^12^. No such increase in calcium was observed in non-transfected HEK cells, or *(-)Therm* transfected HEK cells (Fig. 3), indicating that the magnetic response relies on the thermal activation pathway of TRPV4. We also saw no significant increase in calcium when we applied a constant 275 mT magnetic field for 270 seconds (Fig.3, bottom row), suggesting that the process of magnetization (and not steady magnetic fields) give rise to the calcium signal, which is expected for the magnetocaloric effect.

**Fig. 3.**
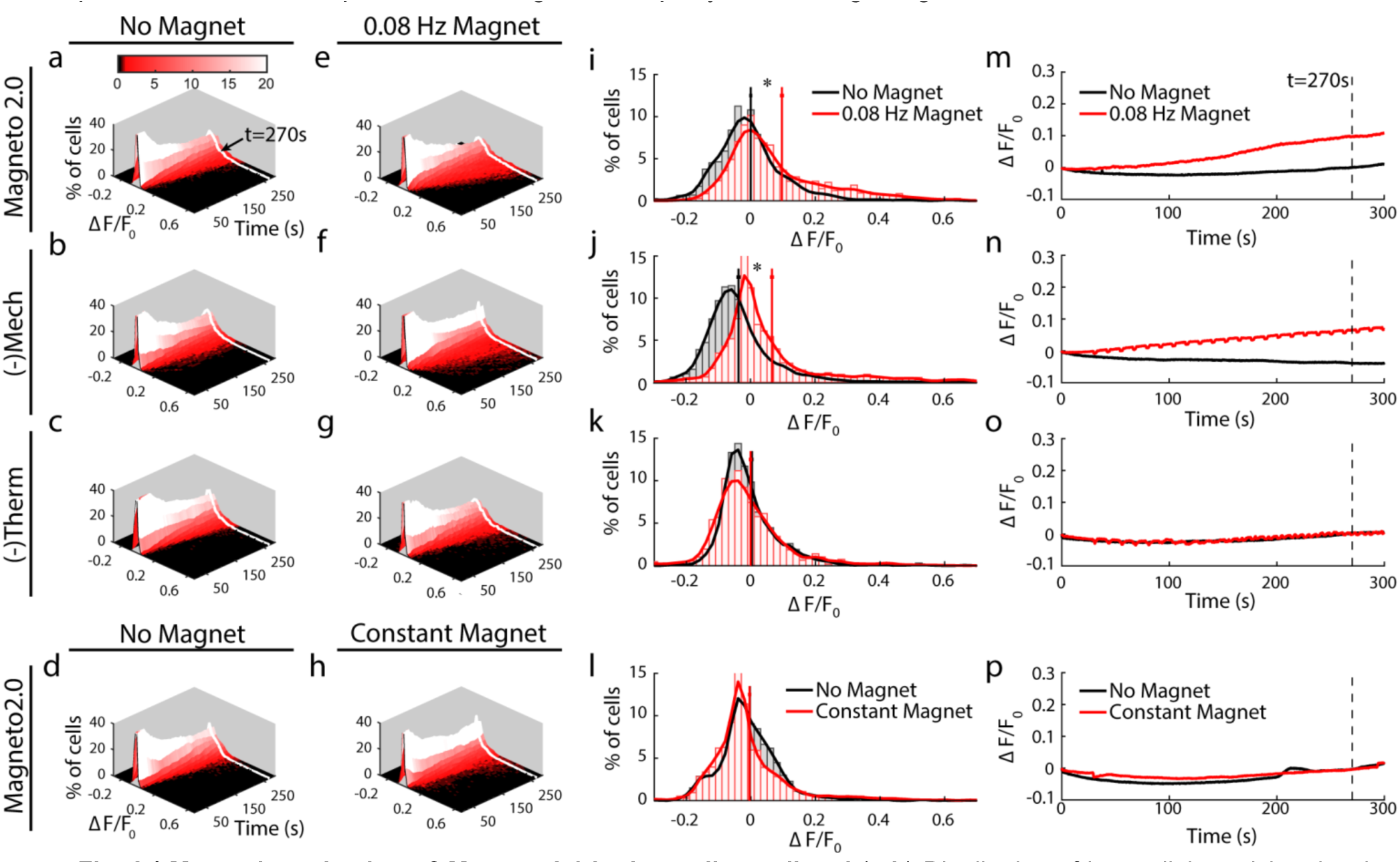
Magnetic activation of *Magneto2.0* is thermally mediated. (**a-h**) Distribution of intracellular calcium levels over time based on calcium sensitive fluorescence imaging (ΔF/F_0_). In the absence of magnetic stimulation (a-d) distribution broadens over time, but remains centered near zero. Under periodic magnetic stimulation (275 mT, 0.08 Hz, beginning at t = 30 s), a small percentage of cells (seen in red tail of the distribution) show an increase in calcium-sensitive fluorescence for *Magneto2.0* (e) and *(-)Mech* (f), but not for *(-)Therm* (g) and *Magneto2.0* under constant magnetic field, shifting the mean of the distribution (h). (**i-l**) Histograms taken from the data in (a-h) show the distribution of fluorescence values at t = 270s with no magnetic stimulation (black) and with magnetic stimulation (red) (bin size 0.2 ΔF/F). These histograms correspond to the white lines in (a-h). Vertical red and black lines represent the mean value of these distributions with and without magnetic stimulation, respectively. Error bars show the s.e.m. for each histogram. (**m-p**) Plotting the mean value of ΔF/F_0_ over time shows that *Magneto2.0* and *(-)Mech* both show statistically significant responses to magnetic stimulation, while *(-)Therm* and *Magneto2.0* in a constant magnetic field do not. Shaded regions are s.e.m. and the vertical dashed line marks t = 270s. Mean ΔF/F_0_ values are averaged using a 20 s sliding window. Equal numbers of “No Stimulation” and “Magnetic Stimulation” experiments were performed for each condition. The total number of cells measured from separate cell cultures are (indicated as total number of cell/number of separate cell cultures): *Magneto2.0*, *n* = 1573/14 (no stimulation), *n* = 1510/14 (magnetic stimulation); *(-) Mech*, *n* = 1290/11 (no stimulation), *n* = 1587/11 (magnetic stimulation); *(-) Therm*, *n* = 1724/10 (no stimulation), *n* = 1073/10 (magnetic stimulation). Statistical significance was measured 270 s after the start of the experiment using a left tailed Wilcoxon, (*Magneto2.0*: P < 0.001; (-) Mech: P < 0.001; (-) Therm: P = 0.96; Cst Mag: P = 1.0) s.e.m. is calculated using n = number of cells. Average ΔF/F_0_ for individual recordings are plotted in Fig. S4.

Our calculations and experimental results support the magnetocaloric hypothesis for magnetic activation of TRPV4-ferritin fusion proteins, but more work is needed to confirm this activation mechanism. In particular, our model relies on decreased thermal conductance (*g*\*) and local heat absorption (*c*\*) due to the nanoscale separation distance between the ferritin nanoparticle and channel protein. While experimental evidence supports these phenomena in synthetic magnetic nanoparticles, similar experiments with ferritin along with improved theoretical understanding of heat transport at the nanoscale will help constrain the estimates of g*. In addition, better biophysical understanding of the thermal gating mechanisms of TRP channels will further improve our estimates of gating by the magnetocaloric effect. Together, this work will help confirm or disprove the magnetocaloric gating hypothesis. More sensitive magnetogenetic channels will also improve our ability to understand the activation mechanism by enabling more quantitative experiments. For example, single channel electrophysiology would provide a more detailed description of channel activity, but is prohibitively laborious if only a small percentage of channels are activated by the magnetic stimuli. Additionally, stronger calcium or voltage responses would allow researchers to study quantitative differences between stimulation protocols that would help uncover the underlying activation mechanism.

In addition to the experimental evidence supporting a thermal resistance correction factor as high as nine to ten orders of magnitude, we note that such large correction factors are not unprecedented for nanoscale distances. For example, it is a widely accepted that the Raman signal from molecules within a few nanometers of a metal surface can be increased by 7-14 orders of magnitude^38^. Thus, nanoscale separation distances can produce surprisingly large effects on physical processes.

The giant thermal resistance values (1/g*) required for our theory (and supported by multiple experiments^27,9,26,28,29^) may also have implications for high-frequency magnetic stimulation of ferritin-TRP assemblies. Although the specific absorption rate of ferritin in alternating fields may be too small to produce significant temperature changes in a volume of fluid, the enhanced surface temperature created may produce local temperature changes sufficient to gate nearby thermoreceptors. Thus, the giant thermal resistance could explain the enhanced effect of both magnetocaloric heating and heating due to relaxation losses. It should be noted, however, the magnetocaloric effect may not play a significant role in high-frequency alternating magnetic fields. Because the magnetocaloric effect predicts that particles will heat during magnetization and cool during demagnetization, we expect the magnetocaloric effect to produce no net temperature change in a rapidly oscillating magnetic field.

In the case of a slowly varying field, a nonlinear response of the cell and/or channel is required to produce a net physiological change due to the periodic heating and cooling produced by the magnet. For example, during cycles of magnetization, the calcium influx produced by heating TRPV4 must be larger than the net calcium efflux produced during cycles of demagnetization that cool TRPV4. We expect that three mechanisms might contribute to asymmetric responses: i) Calcium is known to modulate TRPV4 activity and thus calcium influx could be amplified by positive feedback on the channel ^39^, ii) Secondary messengers and/or calcium itself can trigger the release of calcium from intracellular calcium stores ^40,41^ iii) The local depolarization can trigger voltage-gated ion channels in neurons, and to a lesser extent, in non-excitatory cells ^34^. In the case of rapidly switching fields (e.g. hundreds of kHz) we expect that the field switches much faster than these nonlinear effects yielding a negligible physiological response from the magnetocaloric effect.

Perhaps the most exciting outcome of the magnetocaloric hypothesis is a rational approach to improve the magnetic response. For example, we predict that improving the heat transfer efficiency or the thermal sensitivity of *Magneto2.0* will improve the magnetic sensitivity. Thus, the magnetocaloric hypothesis provides both a potential explanation for the recently reported magnetogenetic proteins and an approach for developing new, more sensitive constructs that respond to low frequency magnetic stimuli.

## Methods

### Cell culture and molecular biology

HEK293 cells obtained from ATCC were cultured in DMEM (Lonza) supplemented with 10% FBS (Gibco, Lot#1750106) and 1% pen-strep (Lonza). Cells were transfected with pcDNA3.1-*Magneto2.0*-P2A-RFP 4 days prior to recording, using Lipofectamine (Invitrogen) following manufacturer’s recommendations. Cells were replated on sterile coverslips 48 hours before recording, to obtain a confluency of 60-80%. WT *Magneto2.0* was obtained from AddGene and mutations were made using the Q5 Site-Directed Mutagenesis Kit from New England Biolab.

### Electrophysiology

The cells were placed in electrophysiology extracellular buffer (eECB, in mM: 145 NaCl, 5 KCl, 3 MgCl_2_, 10 HEPES and 1 CaCl_2_; pH 7.2; adjusted to 320 mOsm with sucrose). Glass patch pipettes with a resistance of 3 to 5 MΩ were filled with intracellular buffer (in mM: 140 KCl, 10 HEPES and 0.5 EGTA; pH 7.2; adjusted to 320 mOsm with sucrose) and brought into contact with the cell membrane to generate seals ≥ 1 GΩ. A negative pressure of -70 mmHg was applied inside the pipettes to gain access to the whole cell configuration. An Axopatch 700A amplifier was used to monitor currents under voltage clamp conditions. The current was filtered at 10 kHz and digitized at 2 kHz using a Digidata 1550 (Molecular Device).

### Calcium Imaging

All calcium recordings were performed in an imaging extracellular buffer (iECB, in mM: NaCl 119, KCl 5, Hepes 10, CaCl_2_ 2, MgCl_2_ 1; pH 7.2; 320mOsm). Cells are incubated with 2 μM Fluo-4 AM (Thermo Fisher Scientific) in culture media for 30 minutes, and rinsed in DMEM for 15 minutes. The coverslip with the cells is then transferred to the recording chamber, covered with iECB and equilibrated at RT for 10 minutes prior to recording. Cells with Fluo-4 were imaged on a Nikon Eclipse inverted microscope with a 20X objective (Nikon S Fluor, N.A.= 0.75; W.D.= 1 mm). For fluorescence excitation, we used an LED with a center wavelength of 470 nm (ThorLabs M470L3). The LED output intensity was set to 160 mW, and filtered to 3% transmittance with ND filters. Images were collected with a Zyla sCMOS Camera (Andor) through a GFP Filter Cube Set (Nikon) and analyzed with Matlab.

### Magnetic stimulation

The magnetic stimulation was delivered by a 1” × 1” cylindrical neodymium rare earth permanent magnet (grade N48, Apex Magnet) on a computer-controlled translation stage (Thorlabs). To collect a baseline fluorescence value, no magnetic stimulation was performed for the first 30 seconds of imaging. After the initial 30 seconds of imaging, the magnet was brought within approximately 8 mm of the coverslip at a frequency of 0.08 Hz. At that distance, the magnetic field is predicted to be 275 mT based on manufacturer’s specifications, and measured in excess of 200 mT (GM-2 gaussmeter, AlphaLab Inc.). The periodic magnetic stimulation was applied for 270 s and the imaging and magnet movements were synchronized using Axopatch (Molecular Device). For each coverslip, a recording was first performed in the absence of magnetic stimulation (“No Stim”), the microscope was then moved to a different field of view (FOV) for magnetic stimulation (“0.08 Hz Stim”). This approach ensured that for each experiment, the cells were exposed to the same illumination conditions and exposed only once to the magnetic stimulation protocol. After magnetic stimulation, the coverslip was discarded. The experiments were performed at 23-25 °C and recordings occurred within 30 minutes of the cell being removed from the incubator.

### Mechanical and Thermal Stimulation

Mechanical and thermal responses were measured via calcium imaging of cells under constant fluid flow in a microfluidic chamber. The recording chamber consisted of a central chamber (~100 μL), three inlet ports, and one outlet port. Coverslips with adherent cells were placed into the chamber, and a PDMS lid provided a watertight seal and thermal insulation during perfusion. The three inlet ports were connected to valve-controlled reservoirs, allowing a gravity-driven exchange of the buffer at 2 mL/min. For each coverslip, calcium activity was monitored during the perfusion of 320 mOsm iECB at 23 °C for 30 s. 240 mOsm iECB (*mechanical stimulation*) or heated 320 mOsm iECB (*thermal stimulation*) were then perfused for 60 s, followed by a return to 320 mOsm iECB at 23 °C for 30 s. For thermal stimulation iECB was heated with an in-line heater (Warner Instrument) to yield a temperature of 40 °C in the recording chamber (measured via thermocouple). Upon perfusion of heated iECB, a small decrease in Fluo-4 intensity is consistently observed in all samples. This stimulation artifact is believed to be due to a temperature-dependent Fluo-4 extrusion ^42^ or decrease in F_max_ ^43^.

### Image processing and analysis

Calcium data was analyzed using custom algorithms developed in MATLAB (MathWorks). First, transfected cells were identified based on mCherry expression, and regions of interest (ROIs) corresponding to individual transfected cells were automatically selected via our segmentation algorithm. We then calculated the percent change in fluorescence (ΔF/F_0_) for each ROI based on the average fluorescence value divided by the average fluorescence value of the first captured image, F_0_. Rarely, sample movement or focal shifts would accompany magnet movement resulting in large periodic artifacts in the imaging data. These data sets were discarded with the exception of *(-)Therm*, where the motion artifacts were small compared to the magnetic field induced changes in fluorescence.

## Acknowledgements

We thank Caleb Kemere, Joff Silberg, Ashley Benham, Douglas Natelson, Polina Anikeeva, Ali Güler, Cecilia Clementi, Mikhail Shapiro, and Markus Meister for their technical assistance or constructive discussions related to this manuscript. We thank Amina Qutub and Arun Mahadevan for their assistance with the image segmentation algorithms.

